# Arabidopsis inositol phosphate kinases, IPK1 and ITPK1, constitute a metabolic pathway in maintaining phosphate homeostasis

**DOI:** 10.1101/270355

**Authors:** Hui-Fen Kuo, Yu-Ying Hsu, Wei-Chi Lin, Kai-Yu Chen, Teun Munnik, Charles A. Brearley, Tzyy-Jen Chiou

**Affiliations:** Agricultural Biotechnology Research Center, Academia Sinica, Taipei 115, Taiwan; Swammerdam Institute for Life Sciences, University of Amsterdam, Science Park 904, 1098XH Amsterdam, The Netherlands; School of Biological Sciences, University of East Anglia, Norwich Research Park, Norwich, Norfolk, NR4 7TJ, U.K.

**Keywords:** Inositol phosphate, inositol pentakisphosphate 2-kinase (IPK1), inositol 1,3,4-trisphosphate 5/6-kinase 1 (ITPK1), phosphate homeostasis, D/L-inositol 45 3,4,5,6-tetrakisphosphate, phosphate starvation response, diphosphoinositol pentakisphosphate, phytate biosynthesis, *Arabidopsis thaliana*

## Abstract

Emerging studies have implicated a close link between inositol phosphate (Ins*P*) metabolism and cellular phosphate (P_i_) homeostasis in eukaryotes; however, whether a common Ins*P* species is deployed as an evolutionarily conserved metabolic messenger to mediate P_i_ signaling remains unknown. Here, using genetics and Ins*P* profiling combined with P_i_ starvation response (PSR) analysis in *Arabidopsis thaliana*, we showed that the kinase activity of inositol pentakisphosphate 2-kinase (IPK1), an enzyme required for phytate (inositol hexakisphosphates; Ins*P*_6_) synthesis, is indispensable for maintaining P_i_ homeostasis under P_i_-replete conditions, and inositol 1,3,4-trisphosphate 5/6-kinase 1 (ITPK1) plays an equivalent role. Although both *ipk1-1* and *itpk1* mutants exhibited decreased levels of Ins*P*_6_ and diphosphoinositol pentakisphosphate (PP-Ins*P*_5_; Ins*P*_7_), disruption of another ITPK family enzyme, ITPK4, which correspondingly caused depletion of Ins*P*_6_ and Ins*P*_7_, did not display similar P_i_-related phenotypes, which precludes these Ins*P* species as effectors. Notably, the level of D/L-Ins(3,4,5,6)*P*_4_ was concurrently elevated in both *ipk1-1* and *itpk1* mutants, which implies a potential role for Ins*P*_4_ in regulating P_i_ homeostasis. However, the level of D/L-Ins(3,4,5,6)*P*_4_ is not responsive to P_i_ starvation that instead manifests a shoot-specific increase in Ins*P*_7_ level. This study demonstrates a more nuanced picture of intersection of Ins*P* metabolism and P_i_ homeostasis and PSR than has previously been elaborated, and additionally establishes intermediate steps to phytate biosynthesis in plant vegetative tissues.

**Significance Statement:** Regulation of phosphate homeostasis and adaptive responses to phosphate limitation is critical for plant growth and crop yield. Accumulating studies implicate inositol phosphates as regulators of phosphate homeostasis in eukaryotes; however, the relationship between inositol phosphate metabolism and phosphate signaling in plants remain elusive. This study dissected the step where inositol phosphate metabolism intersects with phosphate homeostasis regulation and phosphate starvation responses.

## Introduction

Elemental phosphorous (P) in its oxidized form, phosphate (PO_4_^3−^; P_i_), is essential to all life. As a component of nucleic acids, proteins, phospholipids and numerous intermediary metabolites, P_i_ is key to energy metabolism and signal transduction. Plants preferentially acquire P in the form of P_i_ from the rhizosphere, where P_i_ is often limiting owing to its sorption to soil particles and leaching (Holford, 1997). As an adaptation to fluctuating external P_i_ concentrations, plants have evolved intricate regulatory mechanisms to maintain cellular P_i_ homeostasis in vegetative tissue in order to coordinate growth, development, and reproduction, whereas in seeds, P_i_ is reserved in phytate (inositol hexakisphosphate, Ins*P*_6_) that accumulates to several percentage dry weight (Raboy, 1997). In response to P_i_ deficiency, plants initiate a systematic response, termed the P_i_-starvation response (PSR), which involves transcriptional, metabolic, and morphological reprogramming, to enhance P_i_ uptake, allocation, remobilization, and conservation (Rouached et al., 2010; Yang and Finnegan, 2010). Under P_i_-replete or -replenishment conditions, plant cells relieve PSR and store excess P_i_ in the vacuole to avoid cellular toxicity as a result of cytosolic P_i_ surge (Müller et al., 2004; Lin et al., 2013; Liu et al., 2015; Liu et al., 2016). How plant cells perceive external and cellular P_i_ status to maintain P_i_ homeostasis remains elusive despite reports of multiple factors proposed to be signaling molecules, including sugar, phytohormones, microRNAs, Ins*P*s and P_i_ *per se* (Martin et al., 2000; Franco-Zorrilla et al., 2005; Liu et al., 2005; Bari et al., 2006; Chiou et al., 2006; Chiou and Lin, 2011; Puga et al., 2014; Wang et al., 2014).

Inositol phosphates (Ins*P*s) are metabolites of variable phosphorylation on a carbohydrate core, inositol, and are present in all eukaryotes. They are synthesized by evolutionarily conserved enzymes (Irvine and Schell, 2001) and play important roles in diverse cellular processes by functioning as structural and functional cofactors, regulators, and second messengers (Shears et al., 2012). According to the definition of a ‘signal’, being that of agonist-responsive change in concentration that is recognized by a defined receptor (Shears et al., 2012), only very few Ins*P*s can be considered true signaling molecules, including Ins(1,4,5)*P*_3_ in the context of Ca^2+^ signaling (Berridge, 2009) and Ins(3,4,5,6)*P*_4_ as a regulator of the conductance of the Ca^2+^-activated chloride channels (Vajanaphanich et al., 1994; Shears et al., 2012). In plants, Ins*P*s have been hypothesized to mediate signaling of multiple physiological processes, including stomatal closure, gravitropism, drought tolerance, and defense (Lemtiri-Chlieh et al., 2000; Lemtiri-Chlieh et al., 2003; Perera et al., 2006; Mosblech et al., 2008; Murphy et al., 2008; Perera et al., 2008; Laha et al., 2015); however, their roles as signaling messengers in most cases have not been assessed extensively.

The first elaboration of the involvement of Ins*P*s in eukaryotic P_i_ homeostasis was revealed when a rabbit cDNA clone was shown to stimulate P_i_ uptake when ectopically expressed in Xenopus oocytes (Norbis et al., 1997). This so-called P_i_ uptake stimulator (P_i_US) was identified to encode an Ins*P*_6_ kinase (IP6K) that converts Ins*P*_6_ to diphosphoinositol pentakisphosphates (PP-Ins*P*_5_ or Ins*P*_7_) (Norbis et al., 1997; Schell et al., 1999). In yeast, disruption of multiple enzymes responsible for biosynthesis of Ins*P*s and diphosphoinositol phosphates (PP-Ins*P*s) (e.g., Plc1p, Arg82p, and Kcs1p) led to constitutive activation of a P_i_ starvation-responsive phosphatase-coding gene, Pho5, under P_i_-replete conditions (Auesukaree et al., 2005). Subsequent work showed that the synthesis of Ins*P*_7_ by the other family of PP-Ins*P* kinases (Vip1/PPIP5K), Vip1, is stimulated by P_i_ starvation (Lee et al., 2007) and Ins*P*_7_ binds to Pho81, causing inhibition of the Pho80-Pho85 cyclin-cyclin–dependent kinase complex and unphosphorylation of the Pho4 transcription factor. The resulting reduction in phosphorylation of Pho4 localizes this protein to the nucleus, where it activates P_i_ starvation-inducible genes (Lee et al., 2007; Lee et al., 2008). The synthesis of PP-Ins*P*s is also metabolically linked to the synthesis of the main intracellular P_i_ storage molecule, a linear chain of polyphosphate (polyP), and the yeast IP6K mutant, *kcs1*Δ, fails to accumulate polyP (Auesukaree et al., 2005; Lonetti et al., 2011).

Cellular adenylate energy is influenced by P_i_ availability and PP-Ins*P* synthesis (Boer et al., 2010; Szijgyarto et al., 2011; Choi et al., 2017) and itself regulates the synthesis of PP-Ins*P* (Voglmaier et al., 1996; Saiardi et al., 1999; Wundenberg et al., 2014). Together with the genetic and molecular evidence described previously, PP-Ins*P*s have been proposed as metabolic messengers that mediate P_i_ signaling. This hypothesis is further supported by structural and biochemical analyses demonstrating that Ins*P*s and PP-Ins*P*s bind to an evolutionarily conserved SYG1/PHO81/XPR1 (SPX) domain present in proteins that play key roles in P_i_ sensing and transport, with PP-Ins*P*s showing the highest binding affinity (at sub-micromolar concentrations for yeast and animal protein) (Secco et al., 2012; Secco et al., 2012; Wild et al., 2016). Disruption of Ins*P*/PP-Ins*P* binding sites in the SPX domain impaired yeast vacuolar transporter chaperone (VTC)-dependent polyP synthesis and failed to complement P_i_-related phenotypes of the Arabidopsis *phosphate 1* (*pho1*) mutant (Wild et al., 2016). Despite the wealth of current investigation, the evidence for PP-Ins*P*s as evolutionally conserved messengers in eukaryotic P_i_ signaling is scattered, confounded by the absence of Pho80-Pho85-Pho81 homologs in other eukaryotic organisms and the contradictory responses of Ins*P*_7_ levels to P_i_ starvation reported in yeast (Lee et al., 2007; Wild et al., 2016) as well as the presence of a Vip1-independent PHO signaling pathway (Choi et al., 2017).

In plants, a contemporary implication of Ins*P* metabolism in regulation of P_i_ homeostasis comes from a study in which genetic disruption of the kinase responsible for Ins*P*_6_ synthesis, inositol pentakisphosphate 2-kinase (IPK1), causes excessive P_i_ accumulation (Stevenson-Paulik et al., 2005) as a result of elevated P_i_ uptake/allocation activities and activation of a subset of P_i_ starvation-responsive genes (PSR genes) under P_i_-replete conditions (Kuo et al., 2014). In addition to decreased Ins*P*_6_ level, *ipk1* mutation causes a significant change in Ins*P* composition, including accumulation of lower phosphorylated Ins*P* species (e.g., Ins*P*_3_, Ins*P*_4_ and Ins*P*_5_) and decreased levels of PP-Ins*P*s [Ins*P*_7_ and Ins*P*_8_ (*bis*diphosphoinositol tetrakisphosphate)] (Stevenson-Paulik et al., 2005; Laha et al., 2015). The mechanism of IPK1 modulating P_i_ homeostasis and whether Ins*P*s play a role in P_i_-starvation signaling in plants is currently unknown.

As compared with the situation in other eukaryotic organisms, the investigation of biosynthesis of Ins*P*s and their composition in the vegetative tissues of plants is necessarily more complicated than in other eukaryotes due to the presence of complex gene families of Ins*P* biosynthesis enzymes. Mammalian Ins*P* metabolism is dominated by receptor-coupled activation of phospholipase C (PLC) and subsequent metabolic conversion of Ins(1,4,5)*P*_3_ to multiple higher and lower Ins*P*s (Irvine and Schell, 2001), but few plant studies offer detailed identification of Ins*P* species in vegetative tissues due to the limited levels of labeling achieved with *myo*-[^3^H]inositol. Nevertheless, specific short-term non-equilibrium labeling with [^32^P]P_i_ has afforded a metabolic test capable of distinguishing the order in which phosphates are added to the inositol core (Stephens and Downes, 1990; Stephens and Irvine, 1990; Whiteford et al., 1997) and applied to vegetative tissues of plants that revealed a ‘lipid-independent’ pathway of Ins*P*_6_ synthesis (Brearley and Hanke, 1993; Brearley et al., 1997).

Here, using reverse genetics and Ins*P* profiling by [^3^H]inositol and [^32^P]P_i_ labelling, we show that maintenance of P_i_ homeostasis in plants under P_i_-replete conditions depends on the kinase activity of IPK1 and an additional inositol 1,3,4-trisphosphate 5/6-kinase ITPK1. Profile comparison of Ins*P*s between *ipk1-1*, *itpk1*, and another mutant defective in Ins*P*_6_ synthesis, *itpk4*, reveals a correlation between elevated D/L-Ins(3,4,5,6)*P*_4_ [Ins(1,4,5,6)*P*_4_ and/or Ins(3,4,5,6)*P*_4_] level and activation of P_i_ uptake and PSR gene expression. However, the Ins*P* profile in response to P_i_ starvation is distinct from that of the *ipk1-1* and *itpk1* mutants and marked a shoot-specific increase in Ins*P*_7_ level accompanied by ATP increase. Our study reveals a complex relationship between Ins*P* metabolism and P_i_ homeostasis in plants and identifies ITPK4 as a key enzyme in generating Ins*P*_4_ precursors for phytate biosynthesis.

## Results

### Kinase activity of *IPK1* is required for maintenance of P_i_ homeostasis

We previously demonstrated P_i_ overaccumulation in *ipk1-1* mutants associated with activation of PSR genes involved in P_i_ uptake, allocation, remobilization, and signaling (Kuo et al., 2014). Because Ins*P* kinases have been implicated in transcriptional regulation independent of their catalytic activities (Bosch and Saiardi, 2012; Xu et al., 2013; Xu et al., 2013), we examined whether regulation of P_i_ homeostasis by *IPK1* is kinase-dependent. We constructed two forms of IPK1 bearing mutations in conserved kinase motifs (Stevenson-Paulik et al., 2005) (Figure S1A) at Lys168 (*IPK1*^K168A^) or Asp368 (*IPK1*^D368A^), both shown to cause loss of kinase activity *in vitro* (Gonzalez et al., 2010). The expression of wild-type (WT) IPK1 complemented low Ins*P*_6_ content in *ipk1-1* seeds, whereas Ins*P*_6_ levels in seeds of transgenic lines expressing either of the two point-mutated forms of *IPK1* remained as low as that in *ipk1-1* seeds (Figure 1A). These point-mutated *IPK1* forms were expressed both at the transcriptional and translational levels (Figures 1B and S1B), with subcellular protein localization in the cytosol and nucleus, similar to the WT *IPK1* (Figure S1C). These results indicated that Lys168 and Asp368 are required for kinase activity of *IPK1 in vivo*.

**Figure 1.**
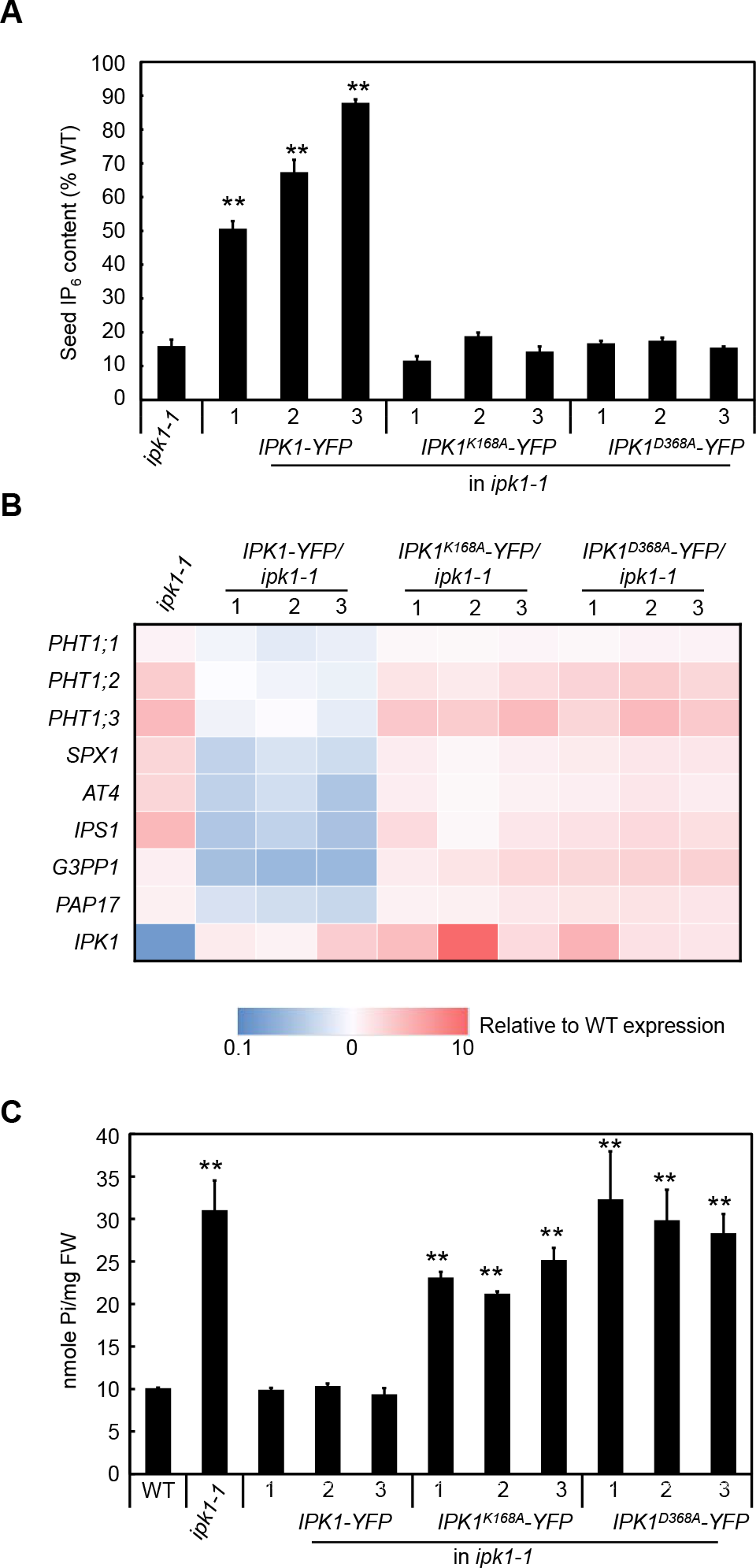
Characterization of kinase-inactive IPK1 transgenic plants. (A) Relative Ins*P*_6_ content (% of WT) in seeds of *ipk1-1* mutants and homozygous transgenic lines expressing C-terminus YFP-tagged wild-type *IPK1* (*IPK1-YFP*), *IPK1*^*K168A*^ (*IPK1*^*K168A*^-*YFP*), or *IPK1*^*D368A*^ (*IPK1*^*D368A*^-*YFP*) coding sequences in the *ipk1-1* mutant background. Error bars, S.E. of n=3-12 independent experiments. (B) Relative expression (to WT) of PSR genes in roots, and (C) P_i_ content in shoots of 14-days after germination (DAG) seedlings. Asterisks indicate significant differences from WT (Student’s *t*-test; **, *P* < 0.005).

In contrast to WT *IPK1*, which was able to restore the P_i_ content of the *ipk1-1* mutant to the WT level, both kinase-inactive *IPK1* forms failed to complement excessive P_i_ accumulation and PSR gene activation in *ipk1-1* (Figure 1B-C). Therefore, the kinase activity of *IPK1* is required for regulation of P_i_ homeostasis. In addition to regulating P_i_ content, the kinase activity of *IPK1* is also required for root system architecture (RSA), because neither of the kinase-inactive IPK1 proteins complemented the PSR-like RSA phenotypes (i.e., reduced primary root and enhanced lateral root growth) of *ipk1-1* (Figure S1D).

### Misregulation of P_i_ homeostasis in *ipk1-1* is not caused by defective Ins*P*_6_-mediated mRNA export

In yeast, Ins*P*_6_ is required for mRNA export by activating the RNA-dependent ATPase activity of DEAD-box protein 5 (Dbp5p) in conjunction with GLFG lethal 1 (Gle1p), and mutations in *ipk1* and *gle1* resulted in mRNA retention in the nucleus and temperature-sensitive growth defects (York et al., 1999; Alcazar-Roman et al., 2006). A conserved mechanism was recently reported in Arabidopsis, and part of the growth defect of *ipk1-1* is attributed to compromised mRNA export due to reduced level of Ins*P*_6_ (Lee et al., 2015). To address whether defective mRNA export in the *ipk1-1* mutant is a cause of the misregulation of P_i_ homeostasis, we examined P_i_-related phenotypes of the mRNA export mutants reported (Lee et al., 2015). As shown in Figure S2, the loss-of-function mutation in the Dbp5 homologous gene *LOW EXPRESSION OF OSMOTICALLY RESPONSIVE GENES 4* (*LOS4*), and inducible *GLE1* RNAi lines exhibited WT P_i_ content (Figure S2A-B) and PSR gene expression (Figure S2C). Furthermore, expression of variants of Gle1 (IS1 and IS2), which exhibit increased Ins*P*_6_ sensitivity to LOS4 stimulation and improved growth defects of *ipk1-1* (Lee et al., 2015), did not reduce P_i_ content or suppress PSR gene activation of the *ipk1-1* mutant (Figure S2C-D). These results suggest that misregulation of P_i_ homeostasis in *ipk1-1* is not caused by defective mRNA export due to reduced Ins*P*_6_ level.

### Genetic dissection of the roles for Ins*P* and PP-Ins*P* biosynthesis enzymes in P_i_ homeostasis regulation

The dependence of P_i_ homeostasis on the kinase activity of IPK1 suggested that the PSR activation signal is derived from Ins*P* biosynthesis. To dissect which step(s) of Ins*P* and PP-Ins*P* biosynthesis controls this signal, we examined P_i_-related phenotypes of mutants defective in several Ins*P* and PP-Ins*P* biosynthesis enzymes previously characterized in Arabidopsis, including myo-inositol-3-phosphate synthases (MIPS1-3) (Torabinejad and Gillaspy, 2006), Ins(1,4,5)*P*_3_ 6-/3-kinases (inositol phosphate multikinases; IPK2α and IPK2β) (Stevenson-Paulik et al., 2002), Ins(1,3,4)*P*_3_ 5-/6-kinase enzymes (inositol phosphate tris/tetrakisphosphate kinases; ITPK1-4) (Wilson and Majerus, 1997; Sweetman et al., 2007), PP-Ins*P* synthesizing enzyme PPIP5K (VIP1/VIH2 and VIP2/VIH1) (Desai et al., 2014; Laha et al., 2015), and a mutant of an Ins*P*_6_ transporter, multidrug resistance-associated protein 5 (MRP5) (Nagy et al., 2009). T-DNA insertional mutants were obtained and confirmed by RT-PCR to be null mutants (Table S1, Figure S3A-B).

Morphologically, none of the mutants displayed growth defects as severe as *ipk1-1* (stunted growth and leaf necrosis), although *mips1, itpk1* and *mrp5-2* mutants were smaller than the WT (Figure 2A). The leaf epinasty and PSR-like RSA phenotypic characteristics of *ipk1-1* mutants (Stevenson-Paulik et al., 2005; Kuo et al., 2014) were observed in *itpk1* and *mrp5-2* mutants (Figures 2A and S3C-D) (Kuo et al., 2014). Analysis of P_i_ content in the shoot tissues revealed that only *itpk1* accumulated excessive P_i_ comparable to *ipk1-1* (Figure 2B), and this phenotype persisted to the mature stage (Figure S3E). Mild but significantly elevated P_i_ content was observed in *mrp5-2* seedlings but was no longer seen at the mature stage (Figures 2B and S3E). Consistent with the elevated P_i_ content, *itpk1* exhibited elevated uptake of P_i_ activity comparable with that of *ipk1-1*, whereas all other mutants showed WT activities (Figure 2C-F). The excessive P_i_ accumulation in *itpk1* mutants could be restored to the WT level by ectopic expression of a genomic construct of the *ITPK1* sequence (Figure S4A), which confirms a role for ITPK1 in regulating P_i_ homeostasis.

**Figure 2.**
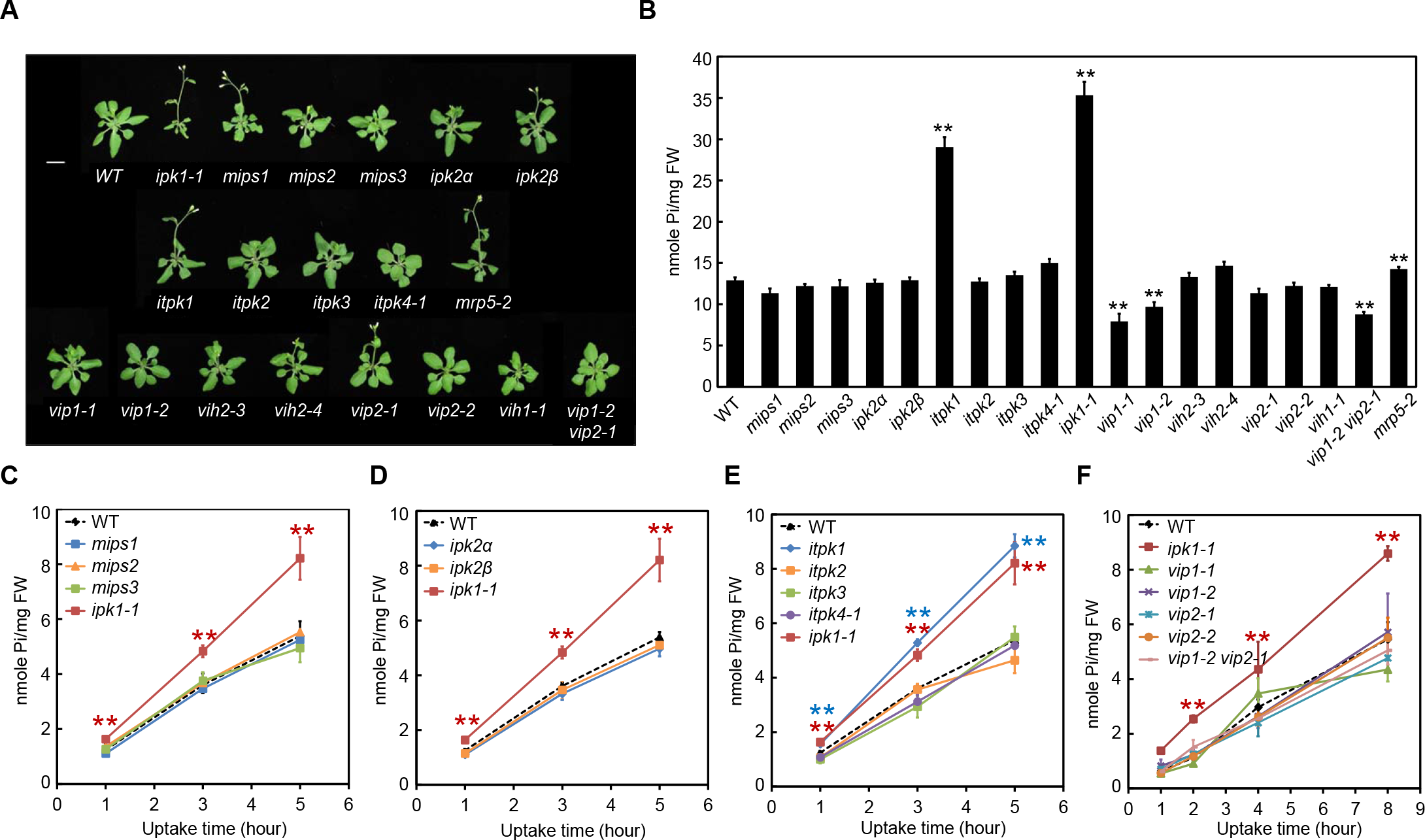
Characterization of mutants defective in Ins*P* biosynthesis enzymes grown under P_i_-replete condition. (A) Morphology of 22-DAG plants grown in P_i_-replete (1 mM) hydroponic medium. Scale bar, 1 cm. (B) P_i_ content in the shoots of 14-DAG seedlings grown on P_i_-replete (1 mM) solid medium. Error bar, S.E. of n=4-21 independent experiments. (C-F) P_i_ uptake activities of 14-DAG seedlings under P_i_-replete (250 μM) growth conditions. Error bars, S.E. of n=3-24 independent experiments. Uptake activities of genotypes in (A-C) were measured in overlapping sets of experiments and plotted separately for clear presentation. Asterisks denote significant differences from the WT (Student’s *t*-test; **, *P* < 0.005).

In addition to decreased Ins*P*_6_ level, levels of Ins*P*_7_ and Ins*P*_8_ are also reduced in *ipk1-1* mutants (Laha et al., 2015). We therefore examined whether PP-Ins*P*s also play a role in the regulation of P_i_ homeostasis or PSR in plants. Two families of kinases, IP6K and Vip/PPIP5K, are involved in PP-Ins*P* synthesis in eukaryotes (Wundenberg et al., 2014); however, only Vip1/PPIP5K homologs are identified in plants and shown to be responsible for Ins*P*_8_ but not Ins*P*_7_ synthesis in Arabidopsis (Mulugu et al., 2007; Desai et al., 2014; Laha et al., 2015). We analyzed mutants defective in each of the two Arabidopsis Vip1/PPIP5K homologs, AtVIP1/VIH2 and AtVIP2/VIH1, and observed slightly decreased P_i_ content in two alleles of *atvip1* mutants (abbreviated as *vip1-1* and *vip1-2*) with T-DNA disrupting the phosphatase-like domain but not in the alleles disrupted in the ATP-grasp kinase domain (*vih2-3* and *vih2-4*) (Figure S3B) (Laha et al., 2015). Three *atvip2* mutants (abbreviated as *vip2-1, vip2-2* and *vih1*) did not show P_i_-content phenotype, but *vip1-2 vip2-1* double mutants exhibited lower P_i_ content comparable to the *vip1-2* single mutant (Figures 2B and S3E), which suggests a dominant role for *vip1* mutation in determining this phenotype. Despite the lower P_i_ content, P_i_ uptake and root-to-shoot allocation activity did not change in the *vip1-1* or *vip1-2* mutants (Figures 2F and S5A). Furthermore, the expression of PSR genes under P_i_-replete conditions and the magnitude of PSR gene activation in response to P_i_ starvation in the *vip1/vih2* and *vip2/vih1* mutants were similar to that in the WT (Figure S5B-C). The cause of reduced P_i_ content observed in *vip1* alleles defective in the phosphatase-like domain is unclear, but the contrasting P_i_-related phenotypes between these *vip1* alleles and *ipk1-1* indicates that the decreased level of Ins*P*_8_ in *ipk1* mutants is not responsible for P_i_ homeostasis misregulation.

### ITPK1 and IPK1 constitute a pathway involved in the maintenance of P_i_ homeostasis

The common phenotypes observed in *itpk1* and *ipk1-1* mutants (i.e., excessive P_i_ accumulation and elevated P_i_ uptake under P_i_-replete growth conditions) suggest that ITPK1 and IPK1 are involved in the same pathway that regulates P_i_ homeostasis.

Consistently, a common set of representative PSR genes was upregulated in *itpk1* and *ipk1-1* mutants (Figure 3A), and overexpression of *ITPK1* or *IPK1* reduced shoot P_i_ content (Figure 3B). Correspondingly, *ITPK1* overexpression significantly decreased P_i_ uptake activity, in contrast to the elevated uptake activity shown by *itpk1* mutants (Figure 3C). In addition, several PSR genes were downregulated in *ITPK1*-overexpressing lines as compared with the WT (Figure 3A, e.g., *PHT1;2*, *SPX1*, *AT4*, *IPS1* and *PAP17*). However, P_i_-uptake activity and PSR gene expression did not differ significantly between IPK1-overexpression lines and the WT (Figure 3A-D).

**Figure 3.**
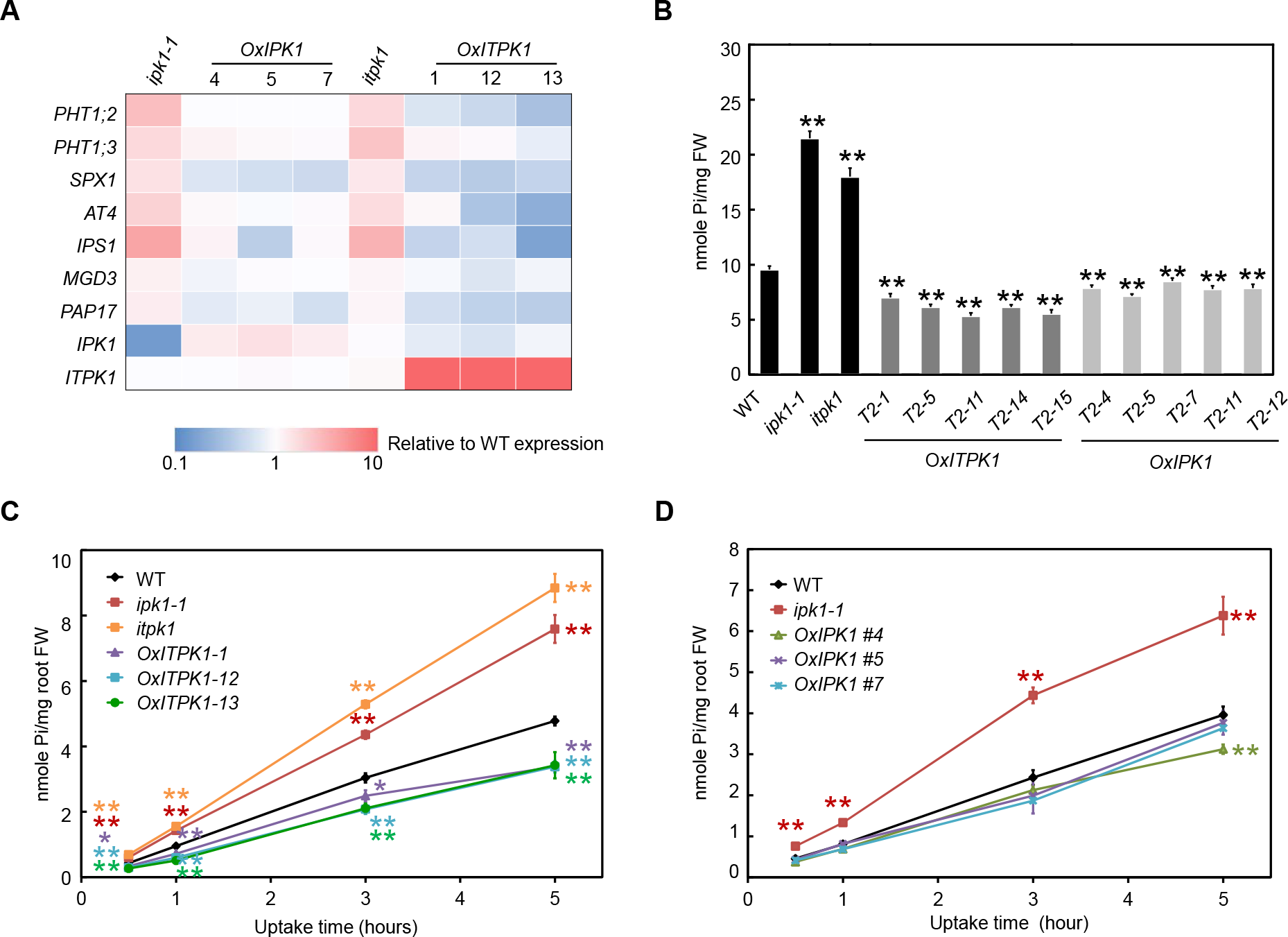
Phenotype similarities between *itpk1, ipk1-1* mutants, and overexpression lines. (A) Relative expression (to WT) of PSR genes in roots of 14-DAG *itpk1*, *ipk1-1*, *IPK1*-overexpression (*OxIPK1*) and ITPK1-overexpression (*OxITPK1*) lines grown under P_i_-replete (1 mM) conditions (see Supporting Table S3 for qPCR raw data and S.E. of 3 independent experiments). Note that qPCR primers for *ITPK1* are located 5’ to the T-DNA insertion site. (B) P_i_ content in shoots of 14-DAG T2 transgenic lines overexpressing *ITPK1* or *IPK1* compared to WT, *itpk1* and *ipk1-1* mutants grown under P_i_-replete (250 μM) condition. Error bars, S.E. of n=6-12 independent experiments. (C and D) P_i_ uptake activities of 14-DAG seedlings grown under P_i_-replete (250 μM P_i_) condition. Error bars, S.E. of n=6-12 independent experiments. Asterisks denote significant differences from the WT (Student’s *t*-test; **, *P* < 0.005).

We drew additional support for the participation of ITPK1 and IPK1 in a common pathway regulating P_i_ homeostasis in terms of their tissue-specific expression patterns and subcellular localization. Promoter-GUS activity assay and RT-PCR analysis demonstrated co-expression of *ITPK1* and *IPK1* throughout development and in specific tissues and cell types, such as vasculature, trichomes and guard cells (Figure 4A-K). In addition, neither gene was transcriptionally responsive to P_i_ status (Figure 4L). The expression of ITPK1 native protein fused to yellow fluorescent protein (YFP), which restored P_i_ content of the *itpk1* mutant to the WT level (Figure S4B), demonstrated co-localization of ITPK1 and IPK1 in the nucleus and cytoplasm (Figures 4M and S1C) (Kuo et al., 2014).

**Figure 4.**
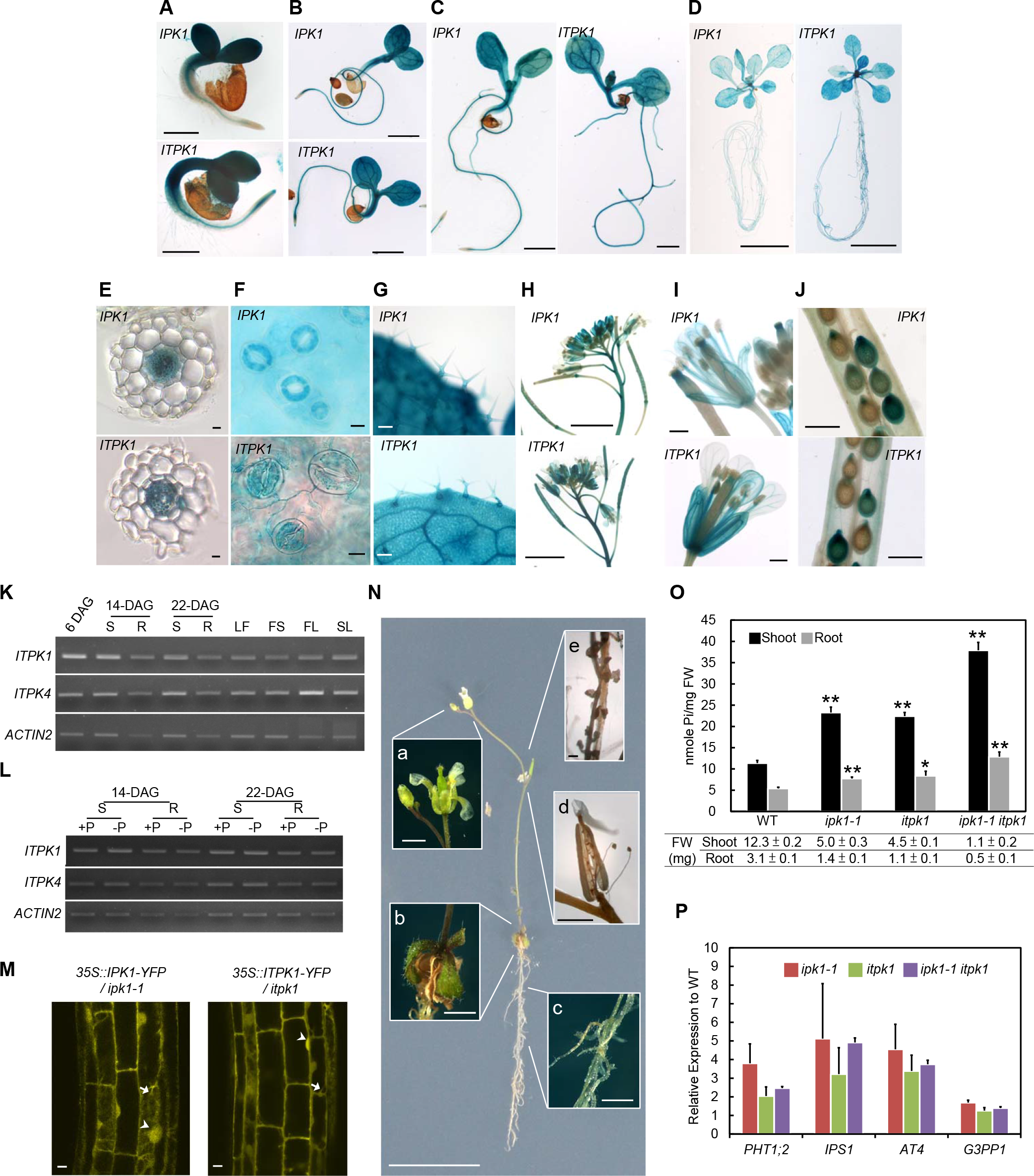
Tissue-specific expression and protein subcellular localization of ITPK1 and IPK1, and phenotypes of *itpk1, ipk1-1* and *itpk1 ipk1-1* double mutants. (A-J) Promoter activities of *IPK1* and *ITPK1* at different developmental stages. (A) 3-DAG; scale bar, 10 μm. (B) 5-DAG; scale bar, 1 mm. (C) 7-DAG; scale bar, 1 mm. (D) 14-DAG; scale bar, 1 cm. (E) Cross section of 14-DAG root; scale bar, 10 μm. (F) Guard cells of 14-DAG leaves; scale bar, 10 μm. (G) Trichome of 14-DAG leaves; scale bar, 0.1 mm. (H) 22-DAG floral tissues; scale bar, 0.5 cm. (I) 22-DAG flowers; scale bar, 0.5 mm. (J) Siliques; scale bar, 0.5 mm. (K and L) RT-PCR analysis of tissue-specific expression of *ITPK1* and *ITPK4* at different developmental stages (K) and in response to P_i_ status (L). S, shoot; R, root; LF, rosette leaves; FS, florescence stem; FL, flower; SL, silique; +P, 250 μM P_i_; −P, 10 μM P_i_. PCR amplification cycles for *ITPK1,* 32; *ITPK4*, 32; *ACTIN2*, 22. (M) Subcellular localization of C-terminus YFP-tagged IPK1 and ITPK1 protein in roots of 10-DAG *ipk1-1* and *itpk1* mutants, respectively; scale bar, 10 μm. Arrows, cytoplasm, arrowheads, nucleus. (N) Morphology of 25-DAG *itpk1 ipk1-1* mutants grown under P_i_-replete (250 μM) conditions. Insets show enlarged images of floral tissues (a), rosette leaves (b), roots (c), mature siliques (d) and aborted seeds (e). Scale bars are 1 cm, 1 mm and 100 μm for the whole plant, insets (a-d) and inset (e), respectively. (O) Tissue-specific P_i_ content and (P) relative expression of PSR genes of 16-DAG seedlings grown on P_i_-replete (250 μM) solid medium. Error bar, S.E. of n=3-6 independent experiments. Asterisks denote significant differences from the WT (Student’s *t*-test; *, *P* < 0.05; **, *P* < 0.005).

We next examined the genetic interaction of *ITPK1* and *IPK1* with a genetic cross between *ipk1-1* and *itpk1* mutants. The *ipk1-1 itpk1* double mutants exhibited more severe growth defects than single mutants (Figure 4N) and those that proceeded to the reproductive stage bore aborted seeds (Figure 4Nd and 4Ne]. Tissue P_i_ content was greater in *ipk1-1 itpk1* double than single mutants, by 50% to 70%, which is likely attributed to the relative 50% to 80% reduction in fresh weight (Figure 4O). Notably, expression of PSR genes in *ipk1-1 itpk1* double and single mutants was comparable (Figure 4P), which suggests IPK1 and ITPK1 function in a common regulatory pathway of P_i_ homeostasis.

### A common elevation of D/L-Ins(3,4,5,6)*P*_4_ in *itpk1* and *ipk1-1* mutants

The observations that maintenance of P_i_ homoeostasis depends on (1) the kinase activity of IPK1, (2) an additional Ins*P* kinase, ITPK1, and (3) the expression level of *ITPK1* and *IPK1* (i.e., contrary P_i_-related phenotypes between mutants and overexpression lines), suggest the contribution of a stoichiometric alteration of Ins*P* metabolites to P_i_ homeostasis regulation. To pinpoint the possible Ins*P* molecules involved in such regulation, we compared Ins*P* profiles of vegetative tissues of the relevant genotypes by *in vivo* labeling with [^32^P]P_i_ and/or *myo*-[^3^H]inositol and HPLC analysis. As shown in Figure 5A, Figure 5B, and *myo*-[^3^H]inositol-labeled chromatogram in Supporting Figure S6A, the *itpk1* mutant shared a significant reduction in Ins*P*_6_ (62 ± 2% WT) with the *ipk1-1* mutant (17 ± 1% WT). To validate that reduced Ins*P*_6_ level is not a cause of misregulated P_i_ homeostasis, with the normal P_i_-related phenotypes exhibited by another low-Ins*P*_6_ mutant *mips1* (Murphy et al., 2008; Kuo et al., 2014), we analyzed the Ins*P* profile of the *mips1* mutant. Unexpectedly, *mips1* mutants exhibited a WT level of Ins*P*_6_ (Figures 5A-B and S6A). For comparison, we also performed profile analysis of other *itpk* mutants and found that two *itpk4* mutants (*itpk4-1* and *itpk4-2;* Table S1) showed a strong reduction in Ins*P*_6_ level comparable to *itpk1* and *ipk1-1* mutants, by 50% and 80%, respectively (Figures 5A-B and S6A). Consistent with the previous report, *itpk4* mutations also significantly reduced Ins*P*_6_ level in seeds, to a similar extent as *ipk1-1* (Figure S7A) (Stevenson-Paulik et al., 2005; Kim and Tai, 2011). The *itpk4* mutants did not show striking morphological phenotypes (Figures 2A and S3C) or P_i_-related phenotypes, such as altered P_i_ content (Figures 2B, S3E and S7B), P_i_ uptake (Figure 2E), or altered PSR gene expression (Figure S7C). RT-PCR and promoter-GUS analysis indicated that *ITPK4* was expressed in the same vegetative tissues as *ITPK1* and *IPK1* (Figure S7D-L), which suggests that ITPK4 is likely involved in the same tissue-specific pool of Ins*P*_6_ biosynthesis. In addition, YFP-tagged ITPK4, which complemented the seed-Ins*P*_6_ phenotype of the *itpk4-1* mutant (Figure S7A), like ITPK1 and IPK1, was also localized to the nuclei and cytoplasm (Figure S7M). Hence, reduced Ins*P*_6_ level alone is insufficient to alter P_i_ homeostasis and ITPK4 is a key enzyme for Ins*P*_6_ biosynthesis in both vegetative tissues and seeds.

**Figure 5.**
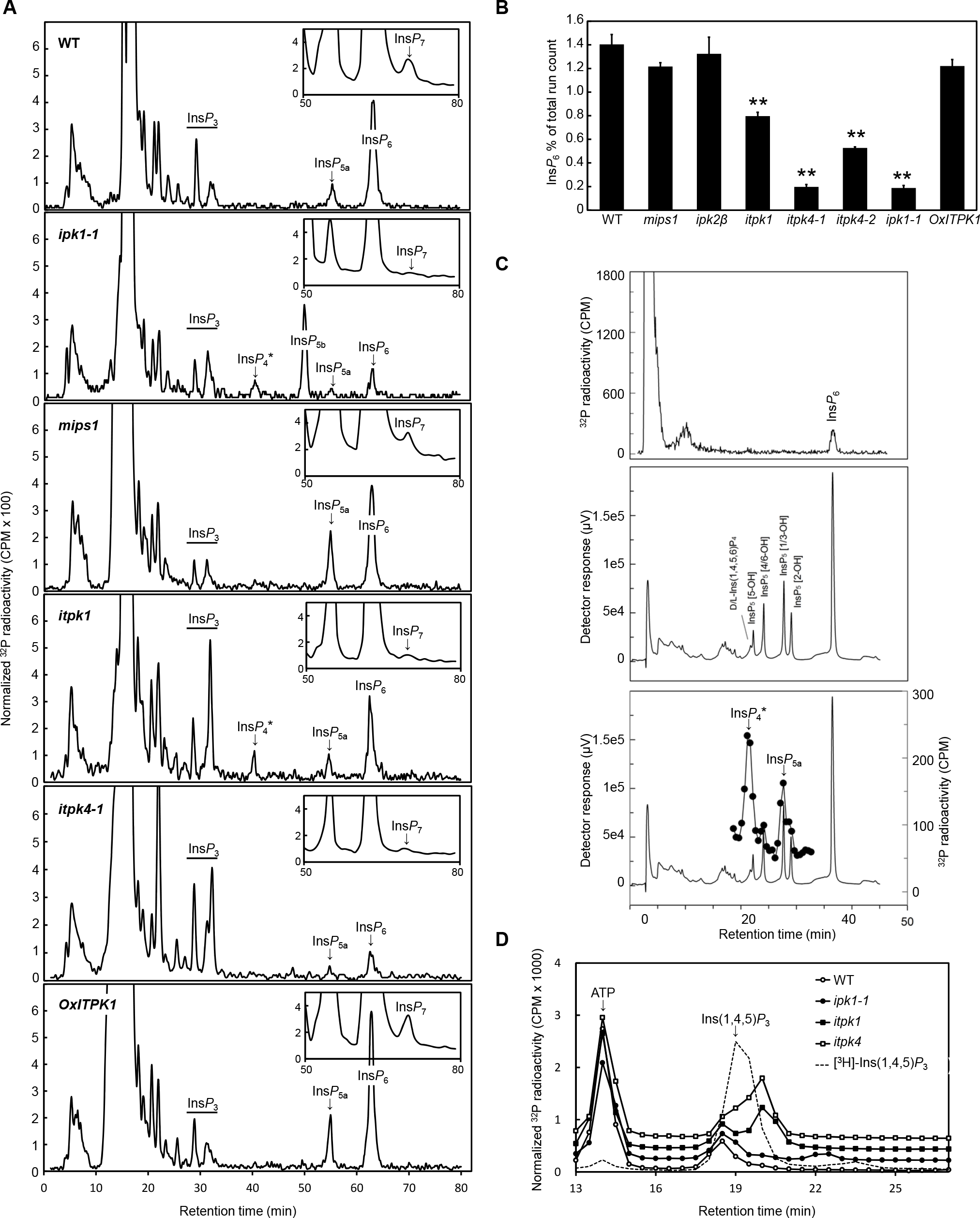
Ins*P* profiles of various genotypes. (A) HPLC analysis of roots extracts from 11-DAG seedlings of various genotypes labeled with [^32^P]P_i_. Ins*P*_5a_, Ins(1,2,4,5,6)*P*_5_ and/or Ins(2,3,4,5,6)*P*_5_ and/or Ins(1,2,3,4,6)*P*_5_ (these three isomers are not resolved on Partisphere SAX HPLC (Brearley and Hanke, 1996); Ins*P*_5b_, Ins(1,3,4,5,6)*P*_5_; Ins*P*_4_*, Ins(1,4,5,6)*P*_4_ and/or Ins(3,4,5,6)*P*_4_; Ins*P*_3_, peaks with the chromatographic mobility of Ins*P*_3s_. Insets show expanded chromatograms of more polar Ins*P*s, obtained by counting 1-min fractions collected from the Flo-Detector eluted from retention time of 50 min onwards. The ordinate is scaled by the same factor for the different genotypes, representing a constant fraction of the largest (P_i_) peak in each chromatogram. (B) Quantification of relative Ins*P*_6_ content (% of total radioactivity per HPLC run recovered in the integrated Ins*P*_6_ peak) in 11-DAG [^32^P]P_i_-labeled seedlings. Error bar, S.E. of n=3-5 independent experiments. Double asterisks denote a significant difference from the WT (Student’s f-test, *P* < 0.005). (C) Identity of Ins*P*_4_* in *itpk1* mutant. An aliquot of extract of [^32^P]P_i_-labeled *itpk1* seedlings (11-DAG) was spiked with a hydrolysate of Ins*P*_6_ and separated on a CarboPac PA200 column with post-column colourimetric detection of Ins*P* peaks as described (Phillippy and Bland, 1988). Upper panel, [^32^P]-radioactivity counted inline on the Flo Detector. Note that Ins*P*_4_ and Ins*P*_5_ are below the level of detection on the Flo Detector, and therefore corresponding fractions (0.5 min) were collected for static counting (lower panel). Middle panel, UV trace obtained from this extract. Ins(1,4,5,6)*P*_4_/Ins(3,4,5,6)*P*_4_ is the latest eluting Ins*P*_4_ on this column and elutes before Ins(1,2,3,4,6)*P*_5_. Lower panel, UV trace overlaid with [^32^P] counts of collected fractions. The retention time of the fractions in the upper and lower panel is corrected for the plumbing delay between the UV detector and the Flo Detector or the fraction collector. The broadness of the [^32^P] peaks (compared to the sharp UV peaks) is a consequence of band broadening after the UV detector (in the Flo Detector and the collected fractions). All the other Ins*P*_4_ isomers elute before 20 min. Ins*P*_5_[5-OH], Ins(1,2,3,4,6)*P*_5_; Ins*P*_5_[4/6-OH], Ins(1,2,3,5,6)*P*_5_ and/or Ins(1,2,3,4,5)*P*_5_; Ins*P*_5_[1/3-OH], Ins(1,2,4,5,6)*P*s and/or Ins(2,3,4,5,6)*P*_5_; Ins*P*_5_[2-OH], Ins(1,3,4,5,6)*P*_5_. Note that a single peak of Ins*P*_4_ was detected in *itpk1* in (A). (D) Isocratic separation and counting of collected fractions for analysis of Ins*P*_3_ isomers. For better presentation, the chromatogram of each genotype was shifted by 200 CPM.

In accordance with decreased Ins*P*_6_ level, Ins*P*_7_ level was also decreased in the *ipk1-1* mutant (Figure 5A) (Laha et al., 2015). Similarly, Ins*P*_7_ level was decreased in *itpk1* and *itpk4* mutants (Figure 5A), and therefore we could not draw a correlation between the reduced Ins*P*_7_ level and the P_i_-related phenotypes observed in *ipk1-1* and *itpk1*. The *ipk1-1* mutant shows significant accumulation of Ins(1,3,4,5,6)*P*_5_ along with reduced Ins*P*_6_ level (Ins*P*_5b_ in Figures 5A and S6A) (Stevenson-Paulik et al., 2005), but in contrast, there was no detectable accumulation in the corresponding Ins*P*_5_ in the *itpk1* mutant. This finding suggests that the elevated Ins(1,3,4,5,6)*P*_5_ level in the *ipk1-1* mutant does not explain the misregulation of P_i_ homeostasis.

Notably, the *itpk1* mutant showed elevated level of an Ins*P*_4_ species with identical chromatographic mobility to that in the *ipk1-1* mutant, which is predominantly Ins(3,4,5,6)*P*_4_ (Ins*P*_4_* in Figures 5A and S6A) (Stevenson-Paulik et al., 2005). The Ins*P*_4_ species in the *itpk1* mutant was further analysed by high-resolution HPLC separation and was co-eluted with D/L-Ins(3,4,5,6)*P*_4_ standard (D/L enantiomers are not separable by existing chromatographic technologies) (Figure 5C). In addition to the increase in Ins*P*_4_ level, levels of earlier eluting Ins*P* species were increased in both *itpk1* and *ipk1-1* mutants, which exhibited the chromatographic mobility of Ins*P*_3_ (Figures 5A and S6A). Because there are 20 possible Ins*P*_3_ isomers, being the most difficult Ins*P* to resolve, isocratic HPLC analysis was performed under conditions designed for optimal resolution of these peaks (Wreggett and Irvine, 1989). As shown in Figure 5D, *ipk1-1* and *itpk1* mutations caused accumulation of distinct Ins*P*_3_ isomers that were not detectable in the WT. Inclusion of an internal standard of *myo*-[^3^H]Ins(1,4,5)*P*_3_ revealed that these isomers are not Ins(1,4,5)*P*_3_, which was shown to present only a trivial fraction of Ins*P*_3_ in plant tissues (Brearley and Hanke, 2000). We conclude that the only common change of Ins*P* species associated with the P_i_-related phenotypes of *ipk1-1* and *itpk1* is the elevated D/L-Ins(3,4,5,6)*P*_4_ level.

### P_i_ starvation induced a shoot-specific increase of Ins*P*_7_

To address whether *itpk1* and *ipk1-1* mutants exhibit an Ins*P* profile that shares a common feature with P_i_-starvation responses, we investigated the change in Ins*P* profiles in shoots and roots of WT plants in response to different P_i_-starvation regimes. Ins*P* profiles were analyzed in seedlings subjected to 1- and 3-day P_i_ starvation, when cellular P_i_ concentrations were significantly reduced and PSR genes induced (Figure S8A-B). To avoid biased quantifications of Ins*P*s caused by elevated P_i_-uptake activities during P_i_ starvation, we performed a pulse-chase experiment with seedlings labeled with [^32^P]P_i_ before P_i_ starvation. Tissues were similarly radiolabeled in every pairwise ‘+P’ vs. ‘−P’ treatment, although more [^32^P] was allocated to shoots than roots (Figure 6A-B).

**Figure 6.**
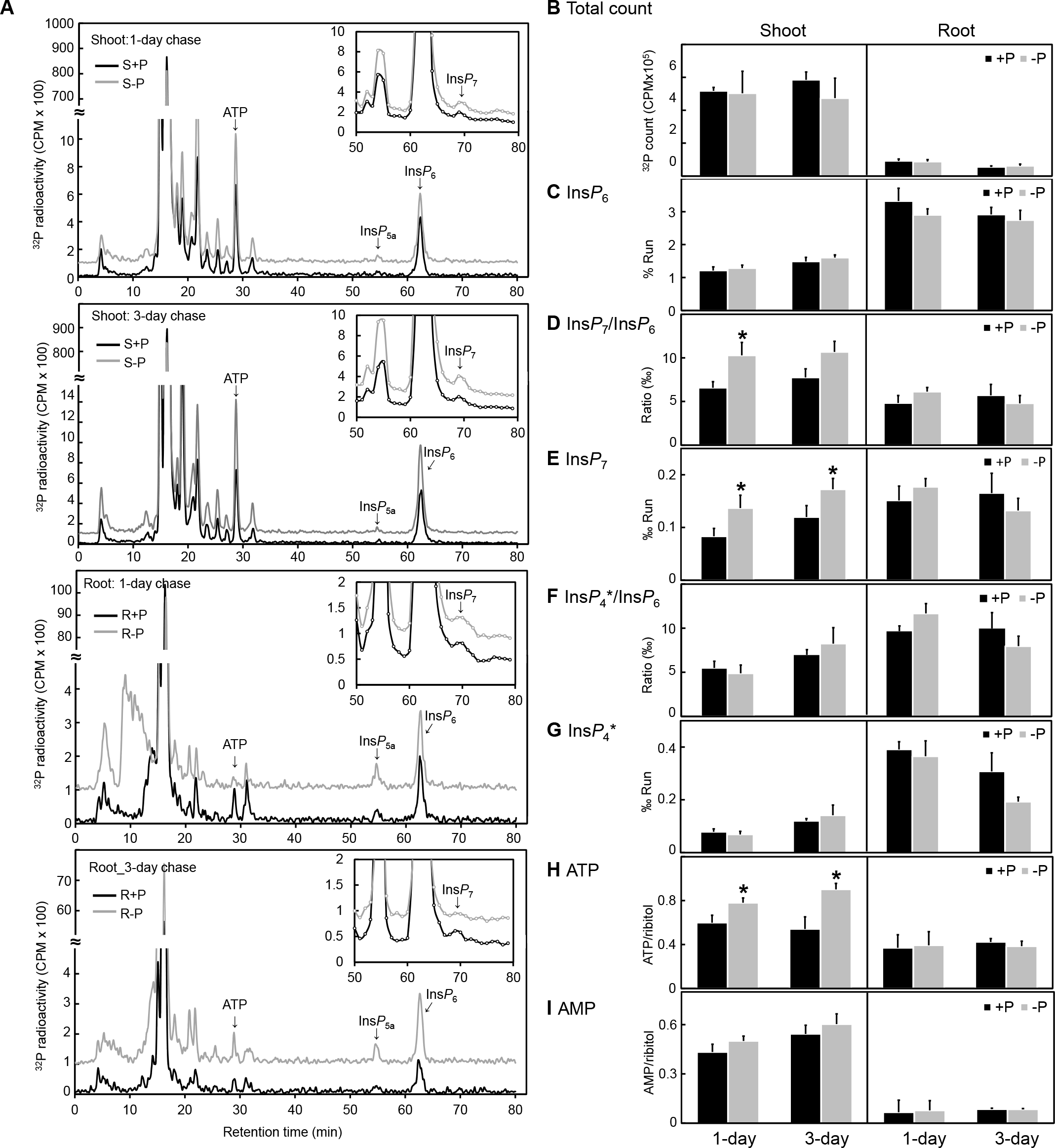
Tissue-specific Ins*P* profiles in response to 1- and 3-day P_i_ starvation. (A) Chromatograms of HPLC analysis of [^32^P]P_i_-labeled WT seedlings after 1- and 3-day P_i_-replete (+P, 250 μM) or P_i_-deficient (−P, 10 μM) treatments. 8-DAG seedlings were labeled with [^32^P]P_i_ under P_i_-replete conditions for 3 days (“pulse“) before transfer to unlabeled media (“chase“). Insets show enlarged chromatograms of more polar Ins*P*s, plotted from scintillation counts of 1-min fractions collected from retention time 50 min onwards. For clearer presentation, [^32^P] radioactivity signals of P_i_ starvation treatment (S-P and R-P) were shifted by 100 CPM. Ins*P*_5a_, Ins(1,2,4,5,6)*P*_5_ and/or Ins(2,3,4,5,6)*P*_5_. (B) Total [^32^P]P_i_ recovered in metabolites of tissues after pulse-chase labeling determined by integration of peaks from in-line flow detection. Error bars, SE of three independently labeled populations of seedlings. (C) Ins*P*_6_ level as percentage of total radioactivity across the gradient recovered in the integrated peak. (D) Ins*P*_7_-to-Ins*P*_6_ ratio determined from counting of fractions in inset (A). (E) Ins*P*_7_ level as per mille total radioactivity derived from Ins*P*_7_-to-Ins*P*_6_ ratio [derived in (D)] multiplied by the Ins*P*_6_ level in (C). (F) Ins*P*_4_*-to-Ins*P*_6_ ratio determined by scintillation counting of fractions as shown in Supporting Figure S9A. Ins*P*_4_* shares common retention time with the elevated peak detected in the *ipk1-1* and *itpk1* mutants (Figure 5). (G) Ins*P*_4_* level as per mille total radioactivity derived from Ins*P*_4_*-to-Ins*P*_6_ ratio (Figure S9A-D) multiplied by the Ins*P*_6_ level in (C). Error bars in (B-G), S.E. of three independent labeling experiments. (H and I) Relative ATP and AMP level per mg fresh weight (FW) derived from normalization to the internal standard ribitol. Error bars, S.E. of n=9 and 7 independent experiments for the shoots and roots, respectively. Asterisks indicate significant differences from the WT (Student’s *t*-test; *, *P* < 0.05).

Overall, the chromatograms did not exhibit prominent profile changes in response to P_i_ starvation in either shoots or roots (Figures 6A and S9). Quantitative analysis indicated no significant change in Ins*P*_6_ level in response to P_i_ starvation in shoots or roots (Figure 6C). Despite no significant change in Ins*P*_7_ level in roots, shoots exhibited mild yet significant increase in Ins*P*_7_-to-Ins*P*_6_ ratio and Ins*P*_7_ level in response to 1- and 3-day P_i_ starvation (Figure 6D-E). We were unable to assess the Ins*P*_8_ level due to the detection limit in our analysis; however, depletion of Ins*P*_8_ caused by *vih2* mutation not affecting the P_i_-starvation response implied that this Ins*P* species does not mediate P_i_ signaling (Figure S5C). Notably, the increase in D/L-Ins(3,4,5,6)*P*_4_ level in *itpk1* and *ipk1-1* mutants was not observed in P_i_-starved WT plants (Figures 6F-G and S9A-D), nor was the level of any Ins*P*_3_ isomer, including Ins(1,4,5)*P*_3_, changed in response to P_i_ starvation (Figure S9E-F).

Because cellular adenylate energy is influenced by P_i_ availability (Boer et al., 2010; Alexova et al., 2017; Choi et al., 2017), and high energy phosphates delivered by ATP are required for pyrophosphorylation (Voglmaier et al., 1996), we examined whether phosphorylated adenine nucleotides are metabolically coordinated with the change in Ins*P*_7_ level in response to P_i_ starvation by LC/MS analysis. ATP increased along with Ins*P*_7_ level specifically in shoots during 1- and 3-day P_i_ starvation, whereas AMP level remained steady (Figure 6H-I), which resulted in a significant increase of ATP/AMP ratio (0.68 ± 0.1 and 1.1 ± 0.1 for 3-day ‘+P’ and ‘−P’ treatment, respectively, *P*=0.009). In conclusion, the changes in Ins*P* profiles of WT seedlings in response to 1- and 3-day P_i_ starvation distinctly differ from those in *itpk1* and *ipk1-1* mutants, which suggests that the mechanism of the ITPK1 and IPK1 contribution to P_i_ homeostasis is distinct from the P_i_-starvation response in WT plants.

## Discussion

In this study, we demonstrated metabolism of distinct Ins*P* species in correlation to P_i_ homeostasis and P_i_ limitation. Under P_i_-replete conditions, the catalytic activity of IPK1 was required for maintenance of P_i_ homeostasis, providing the first evidence of the involvement of Ins*P* metabolism, as opposed to other possible aspects of IPK1 protein function (Figures 1 and S1). This notion is further supported by the identification of an additional Ins*P*-synthesizing enzyme, ITPK1, with a comparable role to IPK1 (Figures 2, 3, S3 and S4). The epistatic relationship of *IPK1* and *ITPK1* in suppressing PSR genes under P_i_-replete conditions, together with their co-expression pattern throughout development and their subcellular co-localization (Figure 4), indicate that ITPK1 and IPK1 constitute an Ins*P* metabolic pathway maintaining P_i_ homeostasis. Ins*P* profiling revealed two distinct common features between *ipk1-1* and *itpk1* mutants: (1) decreases in Ins*P*_6_ and Ins*P*_7_ levels and (2) an increase in D/L-Ins(3,4,5,6)*P*4 level (Figures 5 and S6). In contrast, P_i_ starvation induced a distinct Ins*P* profile from those with *ipk1-1* and *itpk1* mutations (Figure 6), which suggests that *ipk1-1* and *itpk1* mutations affect P_i_ homeostasis by a mechanism other than P_i_-starvation signaling.

### Decrease in Ins*P*_6_, Ins*P*_7_ or Ins*P*_8_ level is not responsible for disturbed P_i_ homeostasis in *ipk1-1* and *itpk1* mutants

The fact that *itpk4* mutants did not exhibit P_i_-related phenotypes comparable to *ipk1-1* and *itpk1* mutants indicates that a decrease in Ins*P*_6_ or Ins*P*_7_ level did not cause the disturbed P_i_ homeostasis under P_i_-replete conditions. The similar tissue/developmental expression pattern and subcellular localization of ITPK4 as ITPK1 and IPK1 suggest that these three enzymes control the same pool of vegetative Ins*P*_6_ and Ins*P*_7_ (Figure S7). While it is possible that radiolabeling does not entirely reflect metabolic (subcellular) pools of different Ins*P* and PP-Ins*P* metabolites, no other methods have been elaborated for measurement of these molecules in plants, never mind their subcellular fractionation. Although we were unable to determine the Ins*P*_8_ level, *vih2* mutants mediating Ins*P*_8_ synthesis *in planta* (Laha et al., 2015) did not phenocopy *ipk1-1* and *itpk1* under P_i_-replete conditions and exhibited normal P_i_-starvation responses (Figures 2, S3 and S5), which suggests that Ins*P*_8_ is unlikely involved in the regulation of P_i_ homeostasis.

We have also ruled out that misregulated P_i_ homeostasis is a secondary consequence of mitigated Ins*P*6-mediated mRNA export by demonstrating that mutations compromising or enhancing Ins*P*_6_-Gle1-Los4 mRNA machinery neither caused comparable P_i_-related phenotypes of *ipk1-1* nor complemented *ipk1-1* (Figure S2). The identification of two *itpk4* alleles with similar reduction in Ins*P*_6_ (and Ins*P*_7_) level in *ipk1-1* and *itpk1*, respectively, without showing P_i_-related phenotypes, also argues against a role for Ins*P*_6_-mediated mRNA export in regulating P_i_ homeostasis (Figures 2, 5 and S7). Of note, although growth retardation of *ipk1-1* is attributed to defective Ins*P*_6_-mediated mRNA export (Lee et al., 2015), *itpk4* mutants did not exhibit growth defects comparable to *ipk1-1* or *itpk1* (Figure 2A). Thus, Ins*P*_6_ reduction may not be the sole cause for the growth defect observed in the *ipk1-1* and *itpk1* mutants.

### Correlation between the increased level of D/L-Ins(3,4,5,6)*P*_4_ and misregulation of P_i_ homeostasis in *ipk1-1* and *itpk1* mutants

Aside from the reduced levels of Ins*P*_6_ and Ins*P*_7_, the most significant common Ins*P* profile change between *itpk1* and *ipk1-1* is the increased accumulation of the Ins*P*_4_ species, shown to predominantly consist of Ins(3,4,5,6)*P*_4_ in the *ipk1-1* mutant (Stevenson-Paulik et al., 2005). The isomeric identity of the Ins*P*_4_ species in the *itpk1* mutant remains to be determined, but human ITPK1 was found a reversible Ins*P* 1-kinase/phosphatase that regulates the level of Ins(3,4,5,6)*P*_4_, an inhibitor of Ca^2^+-activated chloride channels in the plasma membrane (Vajanaphanich et al., 1994; Yang et al., 1999; Ho et al., 2002; Saiardi and Cockcroft, 2008). In tobacco, Ins(3,4,5,6)*P*_4_ is also linked to chloride transport, regulating growth and cell volume in pollen tubes (Zonia et al., 2002). We attempted to test the effect of Ins(1,4,5,6)*P*_4_ or Ins(3,4,5,6)*P*_4_ on P_i_ homeostasis of Arabidopsis seedlings by using membrane-permeant bioactivatable analogues of these two Ins*P* isomers [Bt2-Ins(1,4,5,6)*P*_4_/PM and Bt2-Ins(3,4,5,6)*P*_4_/PM] (Vajanaphanich et al., 1994) but did not observe significant effects on tissue P_i_ accumulation or PSR gene expression. However, the effectiveness of intracellular delivery and metabolism of these Ins*P* analogs on plant tissues remains to be assessed.

In addition to Ins*P*_4_, Ins*P*_3_ showed changes in *ipk1-1* and *itpk1* mutants (Figure 5D). In plants, Ins(1,4,5)*P*_3_ (assayed by a competitive Ins*P*_3_-receptor binding assay) has been linked to several physiological responses, such as gravitropism, salt and drought stresses (Perera et al., 2001; Xiong et al., 2001; Perera et al., 2006; Perera et al., 2008). We demonstrated that neither *ipk1-1* nor *itpk1* mutation affected the levels of Ins(1,4,5)*P*_3_, as measured by radiolabelling approaches. Species that co-elute with this isomer are barely detectable in WT plants (Figure 5D) (Brearley and Hanke, 2000). Because the two mutants showed distinctive Ins*P*_3_ profiles, and neither accumulated Ins(1,4,5)*P*_3_, we did not find any association between changes in specific Ins*P*_3_ and Pi homeostasis.

Because Ins*P* lipids, called polyphosphoinositides (PPIs), also play important roles in cellular signaling and Ins*P* metabolism (Munnik and Vermeer, 2010; Munnik and Nielsen, 2011), we examined whether PPI levels were altered in *ipk1-1* and *itpk1* mutants and found elevated levels of phosphatidylinositol 4,5-bisphosphate [PtdIns(4,5)*P*_2_] in both *ipk1-1* and *itpk1* (Figure S10A-B). We further examined P_i_-related phenotypes in mutants or transgenic lines with elevated levels of PtdIns*P*_2_, i.e., *phosphatidylinositol-phospholipase C2* (*plc2*), *suppressor of actin 9* (*sac9*), and a *PHOSPHATIDYLINOSITOL PHOSPHATE 5-KINASE 3* (*PIP5K3*)-overexpression line (Williams et al., 2005; Kusano et al., 2008; Stenzel et al., 2008; Kanehara et al., 2015). None of these lines were comparable to the *ipk1-1* mutant (Figure S10C-D), which suggests that the increased PtdIns(4,5)*P*_2_ levels in *ipk1-1* and *itpk1* mutants are not likely attributable to the misregulated P_i_ homeostasis.

### P_i_ starvation induced a change in Ins*P* profile distinct from those caused by *itpk1* and *ipk1-1* mutations

Although *ipk1-1* and *itpk1* mutants exhibited characteristic phenotypes of P_i_-starvation responses under P_i_-replete conditions, their Ins*P* profiles were distinct from those under P_i_ starvation, notably the contrasting levels of D/L-Ins(3,4,5,6)*P*_4_, Ins*P*_6_ and Ins*P*_7_ (Figures 5A, 6F-G and S9A-D). The level of D/L-Ins(3,4,5,6)*P*_4_ not being altered by P_i_ starvation suggests these Ins*P* species are not involved in P_i_-starvation signaling in WT plants. The disparate Ins*P* profiles in response to P_i_ starvation versus that caused by *ipk1-1* and *itpk1* mutations imply two distinct P_i_ signaling pathways. In support of this notion, the P_i_-starvation responses persisted in the *ipk1-1* and *itpk1* mutants, in which PSR genes remained inducible under P_i_ starvation (Figure S8C). We observed no distinct alteration of Ins*P* profile in response to P_i_ starvation except for a significant increase in Ins*P*_7_ level of unknown isomeric identity in the shoot of P_i_-starved plants but not in the root (Figure 6D-E), where P_i_-starvation responses also take place. Shoot tissues are more responsive to P_i_ starvation than are roots (Huang et al., 2008; Lin et al., 2008), which has led to a hypothesis that the shoot is the tissue where P_i_ starvation is sensed and the signal initiated (Hammond and White, 2008; Lin et al., 2008). Alternatively, because P_i_ starvation triggers differential transcriptional and metabolic responses between shoots and roots (Wu et al., 2003; Pant et al., 2015), the shoot-specific increase in Ins*P*_7_ level may have tissue-specific physiological significance under P_i_ starvation conditions. It will be important to identify the kinase responsible for Ins*P*_7_ synthesis in plants to address these speculations.

Adenylate energy has been shown to regulate PP-Ins*P*s synthesis, with increased ATP/ADP ratio promoting mammalian IP6K kinase activity (Wundenberg et al., 2014). We observed that the shoot-specific increase in Ins*P*_7_ level was associated with a shoot-specific increase in ATP and ATP/AMP ratio during 1- and 3-day P_i_ starvation (Figure 6H-I). Increases in ATP level in response to P_i_ starvation has been noted in barley leaves (Alexova et al., 2017), which contrasts with the decrease in ATP level during P_i_ starvation reported in yeast (Boer et al., 2010; Choi et al., 2017). P_i_ starvation-induced ATP decreases have been shown in other plant species (Duff et al., 1989; Rao et al., 1989), but concentration ratios of ATP to ADP (or AMP), which control kinetics of cellular metabolism (Pradet and Raymond, 1983), remained unchanged or was increased in those studies. Whether the elevated ATP/AMP ratio drives Ins*P*_7_ accumulation in P_i_-starved shoots awaits further characterization of the Ins*P*_7_ synthesis enzyme. Of note, multiple enzymes involved in adenine nucleotide metabolism have been genetically identified to act upstream of the Pho80/Pho85/Pho81 complex as negative regulators of PHO signaling (Huang and Shea, 2005; Choi et al., 2017). Despite the inter-species difference in strategies for the P_i_-starvation response, accumulating evidence has pointed to a close relationship between adenylate energy status and P_i_ signaling. PP-Ins*P*s are proposed to be ‘metabolic messengers’ that mediate pyrophosphorylation of proteins involved in multiple cellular metabolism, including phosphorylation-based signal transduction pathways in yeast (Saiardi, 2012; Wu et al., 2016). Whether the shoot-specific P_i_ starvation-stimulated Ins*P*_7_ observed in this study has a role in P_i_ signaling by such protein pyrophosphoryaltion remains speculative.

### Significant roles of ITPK family of enzymes in phytate biosynthesis in plant vegetative tissues

Mutation of *IPK1* leads to substantively reduced Ins*P*_6_ level in seeds (Stevenson-Paulik et al., 2005) and vegetative tissues (Stevenson-Paulik et al., 2005; Nagy et al., 2009). The concomitant accumulation of Ins(1,3,4,5,6)*P*_5_ in these tissues/organs (Stevenson-Paulik et al., 2005; Nagy et al., 2009) strongly indicates the dominant contribution of the Ins(1,3,4,5,6)*P*_5_ 2-kinase activity of IPK1 to Ins*P*_6_ synthesis. The coincident accumulation of Ins(3,4,5,6)*P*_4_ in vegetative tissues and seeds (Stevenson-Paulik et al., 2005) may be explained by mass action effects (Hanke et al., 2012), possibly indicating reversibility of the detected Ins(3,4,5,6)*P*_4_ 1-kinase activity (Brearley and Hanke, 2000). The enzyme(s) responsible for producing Ins(3,4,5,6)*P*_4_ in plants are not well defined. In avian erythrocytes, Ins(3,4,5,6)*P*_4_ is the product of 5-phosphorylation of Ins(3,4,6)*P*_3_ and is itself the precursor of Ins(1,3,4,5,6)*P*_5_ (Stephens and Downes, 1990).

In nucleated mammalian cells, the origins of Ins(3,4,5,6)*P*4 have not been tested by the methods of Stephens and Downes (Stephens and Downes, 1990), but the single mammalian ITPK1 is a multifunctional kinase and phosphotransferase that interconverts Ins(3,4,5,6)*P*_4_ and Ins(1,3,4,5,6)*P*_5_ (Chamberlain et al., 2007). The existence in Arabidopsis of a gene family of four inositol tris/tetrakisphosphate kinases (ITPK1-4) complicates study of Ins*P* metabolism. Our identification of significant contributions of ITPK1 and ITPK4 to Ins*P*_6_ synthesis in vegetative tissues focuses attention on the contribution of these enzymes to not just Ins*P*_6_ synthesis but also physiological processes regulated by the intermediate Ins*P*s. *ITPK1* mutation reduces labeling of Ins*P*_6_ by 50%, with concomitant accumulation of D/L-Ins(3,4,5,6)*P*_4_, but because it does so without affecting Ins(1,3,4,5,6)*P*_5_ level (Figures 5A and S6A) suggests that ITPK1 does not likely act as an Ins(1,3,4,5,6)*P*_5_ 1-phosphatase. ITPK1 may be acting at the level of Ins*P*_4_-Ins*P*_5_ interconversion. Remarkably, our studies show that ITPK4, which contributes to nearly 90% of vegetative Ins*P*_6_, and more in seeds, has no effect on the P_i_-starvation response. Our labeling studies showed no increased Ins*P*_4_ accumulation in vegetative tissues (Figures 5A and S6A). This implies that most of the Ins*P*_4_ precursors for Ins*P*_6_ synthesis are generated by this enzyme and the contribution of ITPK4 may lie in its Ins*P*_3_ kinase activity rather than its Ins*P*_4_ isomerase/mutase activity (Sweetman et al., 2007).

### Implication of Ins*P* metabolism in regulating P_i_ homeostasis

Across eukaryotic kingdoms, the SPX domains of a large family of proteins involved in P_i_ sensing and transport have been shown to bind Ins*P*s, thereby regulating SPX-protein activities and their interaction with other proteins (Wild et al., 2016). Although Ins*P*_6_ and PP-Ins*P*s at sub-micromolar concentration exhibited the highest binding affinity to the SPX domains, the lower Ins*P* levels also exhibited physiologically relevant binding affinity at a micromolar range (Wild et al., 2016). Our study has pointed to a significant association between the level of D/L-Ins(3,4,5,6)*P*_4_ and maintenance of P_i_ homeostasis under P_i_-replete conditions but not the P_i_-starvation response. It remains speculative how increases in Ins*P*_4_ level is associated with elevated P_i_ uptake and PSR-gene expression and the future identification of the enantiomerism of D/L-Ins(3,4,5,6)*P*_4_ in the *itpk1* mutant and its interacting protein targets, such as by using Ins*P* affinity screens (Wu et al., 2016), should provide further mechanistic insights. The confounding effects on PHO signaling of Kcs1p (negative) and Vip1p (positive) (Auesukaree et al., 2005; Lee et al., 2007), together with a Vip1-indepdent P_i_-starvation signaling pathway (Choi et al., 2017), suggest the regulatory mechanisms that control P_i_ homeostasis likely involve multiple Ins*P* and PP-Ins*P* species. Different Ins*P* and PP-Ins*P* species may regulate P_i_ homeostasis via their competitive interaction with a spectrum of SPX-domain protein(s). For example, the binding of Ins*P*_6_ and 5-Ins*P*_7_ to OsSPX4/OsPHR2 yielded *K*_d_ of ~ 50 μM and 7 μM respectively (Wild et al., 2016), suggesting that competition between the more abundant Ins*P*_6_ and less abundant PP-Ins*P*s are relevant considerations in SPX function (Wild et al., 2016). Consequently, it will be important to consider the prevailing physiological concentration of potential Ins*P* and PP-Ins*P* competitors. Together with the diverse functions of SPX proteins at different levels of P_i_ homeostasis regulation (Secco et al., 2012; Azevedo and Saiardi, 2017) and our findings presented here, Ins*P*_7_ may not be a general (or conserved) signal, and the role of other Ins*P* intermediates in regulating P_i_ homeostasis need to be considered.

## Experimental procedures

### Plant materials and growth conditions

*Arabidopsis thaliana* mutant lines and their sources are listed in S1 Table; the wild-type line (WT) indicates Col-0 unless specified otherwise. Seeds were surface-sterilized, stratified at 4°C for 1-3 days, and germinated on agar medium of half-strength modified Hoagland nutrient solution containing 250 μM KH_2_PO_4_, 1% sucrose, and 0.8% Bacto agar (Aung et al., 2006). The P_i_-replete (‘+P’) and P_i_-deficient (‘−P’) media were supplemented with 250 μM (or 1 mM as specified) and 10 μM KH_2_PO_4_, respectively. For hydroponic growth, seedlings were germinated and grown on solid media for 10 days before being transferred to half-strength modified Hoagland nutrient solution with sucrose omitted. Plants were grown at 22 °C under a 16-h photoperiod with cool fluorescent white light at 100 to 150 μE m^−2^ s^−1^. For generating *ipk1-1 itpk1* double mutants, both double mutants and isogenic WT progenies were recovered from the F2 population at an equivalent yet lower segregation rate (1%) than expected (6%). Because these two loci are located on different arms of chromosome 5, the reason for this segregation distortion is unknown.

### Measurement of P_i_ content and P_i_ uptake activity

Total P_i_ content and P_i_ uptake activity were measured as described previously (Chiou et al., 2006). To measure the root-to-shoot P_i_ translocation activity, pulse-chase labeling was performed. 14-day after germination seedlings were first incubated in P_i_-replete nutrient solution (half-strength modified Hoagland solution supplemented with 250 μM KH_2_PO_4_) containing ^33^[P]orthophosphate (P_i_) for 3 h (‘pulse’ treatment), then transferred to P_i_-replete nutrient solution without ^33^[P]P_i_ for indicated times (‘chase’ treatment). [^33^P] radioactivity in the plants tissues was measured as the P_i_ uptake assay and the root-to-shoot P_i_ translocation activity was measured by shoot-to-root ratio of ^33^P count.

### Genotype analysis, transgene construction and plant transformation

Primers used for genotyping of T-DNA insertional lines were designed according to SIGnAL (http://signal.salk.edu/tdnaprimers.2.html) and are listed in Supporting Table S2. For constructing kinase-inactive IPK1, nucleotide substitutions were introduced in the primers (5’ phosphorylated; Supporting Table S2) used for PCR amplification by using a vector (pMDC32) containing the IPK1 CDS sequence driven by the 35S promoter as template. PCR product was ligated before transformation and sequences were confirmed before recombination into the Gateway destination vector pK7YWG2.0 (C’-YFP) (Karimi et al., 2007) via LR Clonase enzyme mix (Invitrogen). For complementation analysis, the genomic sequence of ITPK1, including 1 kb upstream of ATG start codon, was amplified by PCR (primers listed in Supporting Table S2) and cloned into pCR8/GW/TOPO (Invitrogen) followed by recombination into the Gateway destination vectors. pMDC99, pMDC32 (Curtis and Grossniklaus, 2003), and pK7YWG2.0 were chosen as destination vectors for complementation, promoter::GUS activity and YFP fluorescence analysis, respectively. All cloned constructs were validated by sequencing analysis before being introduced into Arabidopsis by the floral-dip transformation method (Clough and Bent, 1998).

### RNA isolation, RT-PCR, and qRT-PCR

Total RNA was isolated by using RNAzol reagent (Molecular Research Center) and cDNA was synthesized from 0.5 to 1 μg total RNA by using Moloney Murine Leukemia Virus Reverse Transcriptase (M-MLV RT, Invitrogen) and oligo(dT) primers. Sequences of primers used for RT-PCR and qRT-PCR are in Supporting Table S2. qRT-PCR involved use of the Power SYBR Green PCR Master Mix kit (Applied Biosystems) on a 7500 Real-Time PCR system as instructed. Gene expression was normalized by subtracting the Ct value of *UBQ10* (ΔCt) from that of the gene studied and presented as 2^−ΔCt^. The expression relative to the WT (i.e., fold change relative to the WT) is presented as 2^−ΔΔCt^ (where ΔΔCt =ΔCt-ΔCt^WT^). qPCR raw data is provided in Supporting Table S3.

### GUS staining and fluorescence microscopy

GUS activity of transgenic T2 plants was detected as described (Lin et al., 2005), and the signal was observed under an Olympus SZX12 or a Zeiss AxioSkop microscope. Confocal microscopy images of the YFP signal were obtained by using a Zeiss LSM 510 META NLO DuoScan with LCI Plan-Neofluar x63/1.3 Immersion and Plan-Apochromat ×100/1.4 oil objectives. Excitation/emission wavelengths were 514 nm/520 to 550 nm for YFP.

### Ins*P* profiling of Arabidopsis seedlings and seeds

For Ins*P* profile analysis of Arabidopsis vegetative tissue, seedlings (8-11 DAG) were labelled with *myo*-[2-^3^H]inositol (19.6 Ci mmol^−1^, Perkin Elmer NET114A00; 0.4 mCi mL^−1^ for 5 days) or [^32^P]P_i_ (8500-9120Ci mmol^−1^, Perkin Elmer NEX05300; 0.02 mCi mL^−1^ for 1-3 days accordingly) in half-strength Hoagland’s medium supplemented with P_i_ at levels specified in the text. Ins*P* was extracted from the radiolabeled tissues, roots, shoots or whole seedlings as described (Azevedo and Saiardi, 2006). Extracts were resolved on a 250 × 4.6 mm Whatman Partisphere SAX WVS column fitted with guard cartridge of the same material at a flow rate of 1 mL min^−1^ with a gradient derived from buffer reservoirs containing A, water; B, 1.25M (NH_4_)_2_HPO_4_, adjusted to pH 3.8 with H_3_PO_4_, mixed according to the following gradient: time (min), %B; 0, 0; 5, 0; 65, 100; 75, 100. Isocratic separations of Ins*P*_3_ species were performed at the same flow rate on the same column eluted with 20% buffer B. For *myo*-[^3^H]inositol labeling, fractions were collected every minute from retention time 0 to 30 min and every 0.5 min from 30 min onward, followed by scintillation counting (1:4 ratio column eluent to scintillation cocktail; Perkin-Elmer; ULTIMA-FLO AP). For [^32^P]P_i_ labeling, radioactivity was measured by Cherenkov counting on a Canberra Packard Radiomatic A515 Flow Detector fitted with a 0.5-mL flow cell with an integration interval of 0.1 min (Brearley et al., 1997).

*myo*-[^3^H]inositol and [^32^P]P_i_ exhibited different allocation between tissues *in planta*, with greater [^3^H] labeling of roots (Figure S6B), whereas [^32^P]P_i_ labeled shoots more strongly (Figure 6B). With the exception of experiments to compare the extent of labeling of Ins*P*_6_ between a wide range of genotypes (Figure 5B), performed with whole seedlings, the shoot and root tissues were analyzed independently. Aside from stoichiometric differences of specific Ins*P*s, the Ins*P* profile was in general similar between these two tissues (Figures 6A and S6B).

For analysis of Ins*P*s in seeds, 2 mg seed was homogenized in 500 μl of ice-cold 0.6 N HCl before centrifugation for 15 min to remove cell debris. Aliquots (20 μL) were injected onto a 3 mm i.d. × 200 mm Carbo Pac PA200 HPLC column (Dionex) fitted with a 3 mm × 50 mm guard column of the same material. The column was eluted at a flow rate of 0.4 mL/min with a gradient of methane sulfonic acid (Acros Organics) delivered from buffer reservoirs containing: A, water; B, 600 mM methane sulfonic acid according to the following schedule: time (minutes), % B; 0, 0; 25, 100; 38, 100; 39, 0; 49, 0. The column eluate was mixed by using a mixing tee with a solution of 0.1% w/v ferric nitrate in 2% w/v perchloric acid (Phillippy and Bland, 1988) delivered at a flow rate of 0.2 mL/min, before passage through a 194-uL volume knitted reaction coil (4 m × 0.25 mm i.d.) obtained from Biotech AB, Sweden. The column, mixing tee and reaction coil were held at 35°C. Peaks of Ins*P* were detected at 290 nm with a Jasco UV-2077 Plus UV detector. Chromatographic data were integrated in ChromNav (Jasco) software. The position of elution of different stereoisomers of the different classes of Ins*P*s was determined by the inclusion at regular intervals of a set of standards obtained by extended acid treatment of phytic acid (middle panel in Figure 5C).

### ATP and AMP analysis

Adenylates from plant tissues were extracted as described (Cho et al., 2016). Tissues were homogenized in liquid nitrogen and re-suspended in 2.3% (v/v) TCA containing 200 μg/ml ribitol (250 μl per 100 mg tissue). Homogenates were centrifuged at 13,000 rpm at 4°C for 15 min, and supernatants were recovered and neutralized to pH 6.5-7 by KOH, followed by 30-min incubation on ice. Extracts were centrifuged at 13,000 rpm at 4°C for 15 min and the supernatants were collected for LC/MS quantification with an ultra-performance liquid chromatography (UPLC) system (ACQUITY UPLC, Waters, Millford, MA). The sample was separated with a ZIC-cHILIC column (3-μm particle size, 2.1 × 100 mm, Merck-Millipore). The UPLC system was coupled online to the Waters Xevo TQ-S triple quadruple mass spectrometer. Ribitol was used as internal standard. Characteristic MS transitions were monitored by the negative multiple reaction monitoring (MRM) mode for ATP (m/z, 506→159), AMP (m/z, 346→79), and ribitol (m/z, 151→71). Data acquisition and processing involved use of MassLynx v4.1 and TargetLynx software (Waters Corp.), with intensities of ATP and AMP normalized to ribitol.

## Acknowledgements

We thank Shu-Chen Shen (Confocal Microscope Facility, Scientific Instrument Center, Academia Sinica, Taiwan) for fluorescence microscopy imaging, Chen-Chuan Hsu (Plant Tech Core Facility, ABRC, Academia Sinica, Taiwan) for cross-section imaging of GUS staining in plants, ABRC Metabolomics Core for LC-MS measurement of ATP/AMP, Hsin-Yu Huang, Su-Fen Chiang and Hayley Whitfield (University of East Anglia, Norwich, United Kingdom) for technical support. *vih* and *plc2* mutants were kindly given by Gabriel Schaff (University of Bonn, Germany) and Kazue Kanehara (Institute of Plant and Microbial Biology, Academia Sinica, Taipei, Taiwan), respectively; mutants and transgenic lines of Gle1-Los4 mRNA export machinery were generously provided by Ho-Seok Lee and Hyun-Sook Pai (Department of Systems Biology, Yonsei University, Seoul, Korea). This work was supported by grants from the Ministry of Science and Technology of the Republic of China (MOST 104-2321-B-001-057 and MOST 105-2321-B-001-038 to T.-J. Chiou and postdoctoral fellowship to H.-F. Kuo), the BBSRC grants from the United Kingdom (BB/M022978/1 and BB/N002024/1 to C.A. Brearley), and the Netherlands Organization for Scientific Research (NWO 867.15.020 and 711.017.005 to T. Munnik).

## Author Contributions

H.-F.K. and T.-J.C. conceived the project; H.-F.K., C.B. and T.-J.C. designed the experiments; H.-F.K., Y.-Y.H., W.-C.L., K.-Y.C., T.M. and C.B. performed the research; H.-F.K., C.B., T.M. and T.-J.C. interpreted the results; H.-F.K. and C.B. wrote the manuscript; H.-F.K., T.M., C.B. and T.-J.C. contributed to the final version of this article.

## Supporting Information

Supporting Figure S1. Characterization of kinase-inactive IPK1 transgenic plants

Supporting Figure S2 The role of Gle1-Ins*P*_6_-Los4 mRNA export machinery in *ipk1*-mediated P_i_-related phenotypes

Supporting Figure S3. Genetic and phenotypic characterization of mutants of Ins*P* biosynthesis enzymes

Supporting Figure S4. Complementation of *itpk1* phenotypes

Supporting Figure S5. P_i_ allocation activities and PSR gene expression in the *vip/vih* mutants

Supporting Figure S6. Ins*P* profile of genotypes labeled with *myo*-[^3^H]inositol

Supporting Figure S7. Characterization of *itpk4* mutants and expression pattern of *ITPK4*

Supporting Figure S8. P_i_ starvation responses of WT and various genotypes under different regimes of P_i_ starvation

Supporting Figure S9. Tissue-specific Ins*P* profiles in response to 1- and 3-day P_i_ starvation

Supporting Figure S10. PPI composition in *itpk1, ipk1-1* and *OxITPK1* lines, and P_i_ content in mutants exhibiting elevated PtdIns(4,5)*P*_2_ levels

Supporting Table S1. Mutant lines used in this study

Supporting Table S2. Primers used in this study

Supporting Table S3. RT-qPCR data for gene expression

